# *In situ* and transcriptomic identification of synapse-associated microglia in the developing zebrafish brain

**DOI:** 10.1101/2021.05.08.443268

**Authors:** Nicholas J. Silva, Leah C. Dorman, Ilia D. Vainchtein, Nadine C. Horneck, Anna V. Molofsky

**Affiliations:** Department of Psychiatry and Behavioral Sciences/Weill Institute for Neurosciences, University of California, San Francisco, San Francisco, CA; Neuroscience Graduate Program, Department of Laboratory Medicine, University of California, San Francisco, San Francisco, CA; Kavli Institute for Fundamental Neuroscience, University of California, San Francisco, San Francisco, CA

## Abstract

Microglia are brain resident macrophages that play vital roles in central nervous system (CNS) development, homeostasis, and pathology. Microglia both remodel synapses and engulf apoptotic cell corpses during development, but whether unique molecular programs regulate these distinct phagocytic functions is unknown. Here we identify a molecularly distinct synapse-associated microglial subset in the zebrafish (*Danio rerio*). We found that ramified microglia populated synapse-rich regions of the midbrain and hindbrain between 7 and 28 days post fertilization. In contrast, microglia in the optic tectum were ameboid and clustered around neurogenic zones. Using single-cell mRNA sequencing combined with metadata from regional bulk sequencing, we identified synapse-associated microglia (SAMs) that were highly enriched in the hindbrain, expressed known synapse modulating genes as well as novel candidates, and engulfed synaptic proteins. In contrast, neurogenic-associated microglia (NAMs) were enriched in optic tectum, had active cathepsin activity, and preferentially engulfed neuronal corpses. These data yielded a functionally annotated atlas of zebrafish microglia (https://www.annamolofskylab.org/microglia-sequencing). Furthermore, they reveal that molecularly distinct phagocytic programs mediate synaptic remodeling and cell engulfment, and establish zebrafish hindbrain as a model circuit for investigating microglial-synapse interactions.

## Introduction

Microglia, the dominant immune cells in the central nervous system (CNS), perform critical and diverse functions during brain development and disease, including engulfing synapses, promoting synapse formation, and clearing apoptotic neurons, ^1,2^. However, it is not clear whether these diverse functions are mediated by molecularly distinct subsets of microglia. Single-cell sequencing of mouse microglia reveals transcriptional heterogeneity predominantly during development ^3,4^, whereas functional studies reveal region-specific microglial populations that persist into adulthood ^5–7^. These data suggest that transcriptional profiling in rodents has not yet been able to resolve known functional heterogeneity in microglia. Furthermore, linking these transcriptionally identified subpopulations to functional subsets *in situ* remains challenging. Thus, despite abundant evidence that microglia both modulate synapses and engulf cell corpses during development, the molecular regulation of these functions is not well understood.

The zebrafish (*Danio rerio*) is an increasingly utilized vertebrate model for developmental neuroscience ^8^ that has not been used to study microglial-synapse interactions. Zebrafish contain ontogenetically similar microglia^9–11^, as well as other glial cell types ^12–14^. Fish have a highly homologous immune system to mammals ^15–17^, including meningeal lymphatics ^18^ and a blood brain barrier that matures between 8-10 dpf ^19^. Fish nervous system development also resembles mammals, with the exception that robust neurogenesis persists throughout life, leading to ongoing turnover of neuronal corpses in neurogenic regions, particularly in the midline optic tectum (OT). Much of our functional understanding of fish microglia comes from elegant work in the OT, which has identified molecular mechanisms that drive phagocytosis of neuronal corpses^20–22^. However, in the OT are functionally and ontogenetically distinct from microglia in other CNS regions^23–26^. Furthermore, a distinct subset of microglia in the spinal cord white matter engulfs myelin sheaths^27^, and is linked to leukodystrophy^28^.

These regionally and functionally distinct microglial subsets raise the question of whether synapse-associated microglia also exist in the fish, and if so, whether they play a functional role in CNS development. Bulk transcriptomic sequencing in the fish has begun to uncover key information regarding microglial ontogeny, revealing that microglia populate the CNS in two waves ^29,30^. An embryonic wave derived from the rostral blood island, similar to the yolk sac of mammals, persists for the first two months and is later replaced by an adult population derived from ventral wall of dorsal aorta (VDA) ^29^. Recent bulk sequencing studies describe microglia transcriptional heterogeneity in the adult zebrafish brain, linked to ontogeny, that correlates with differing amounts of bacterial phagocytosis in vitro ^31^. However, most canonical microglial genes identified in mice and humans ^32^ are conserved in both developmental waves of zebrafish microglia ^33–35^. Like in mammals, it is not clear to what extent ontogeny impacts function, thus a focus on linking the functional transcriptome to *in vivo* function is essential.

Here we define a distinct subpopulation of synapse-associated microglia (SAMs) in the juvenile fish using *in situ* characterization as well as single cell and bulk RNA sequencing. SAMs were ramified and expanded in the midbrain and hindbrain after 7 days post fertilization (dpf). These microglia engulf neuronal synaptic proteins as in mammals, and were defined by expression of complement genes (*C1qa, C1qc*) and other novel candidate pathways. In contrast, microglia in the optic tectum clustered near neurogenic regions, were rich in lysosomal gene expression (*Ctsla, Ctsba*), and their phagocytic capacity correlated with functional cathepsin activity. These data define a molecular profile associated with phagocytosis of synaptic proteins and define a novel model system in which to study microglial-synapse interactions.

## Results

### Synapse-associated microglia expand developmentally in the zebrafish hindbrain and are distinct from neurogenic-associated microglia

To characterize microglia in both synaptic and neurogenic regions of the fish brain, we performed immunohistochemistry using the myeloid reporter line *Tg*(*mpeg:EGFP*) ^36^. We used the presynaptic vesicle marker SV2 to demarcate synapse rich regions, and BrdU to identify neurogenic zones. We found that microglia colocalize with synapse rich regions of the midbrain and hindbrain as early as 7 days post fertilization (dpf) and increased in number over the late larval to juvenile period (14-28 dpf; **Fig. 1A, B**). By 28 dpf most microglia in the midbrain and hindbrain were colocalized with the presynaptic marker, SV2 (65% and 80%, respectively; **Fig. 1C,D**). In contrast, >60% of microglia in the optic tectum (OT) clustered near the BrdU+ neurogenic zone (**Fig. 1E,F**), consistent with their known roles in eliminating apoptotic cell corpses ^24,26^. Immunostaining with the commonly used microglia antibody 4C4 were consistent with these findings and indicated that the majority (70-90%) of mpeg:EGFP+ cells in the brain are microglia, with a modestly lower proportion in the midbrain and hindbrain relative to OT (**Fig. S1A-D**). We next characterized microglial morphology in hindbrain and OT. We found that microglia (mpeg:EGFP+4C4+) in the hindbrain were ramified and more closely resembled microglia in the postnatal rodent brain, whereas OT microglia were significantly more ameboid, as quantified by increased sphericity (**Fig. 1G,H**). In summary, we identified a ramified synapse-associated microglia (SAM) enriched in the hindbrain and midbrain that was distinct from amoeboid neurogenic-associated microglia (NAM) in the optic tectum.

**Figure 1.**
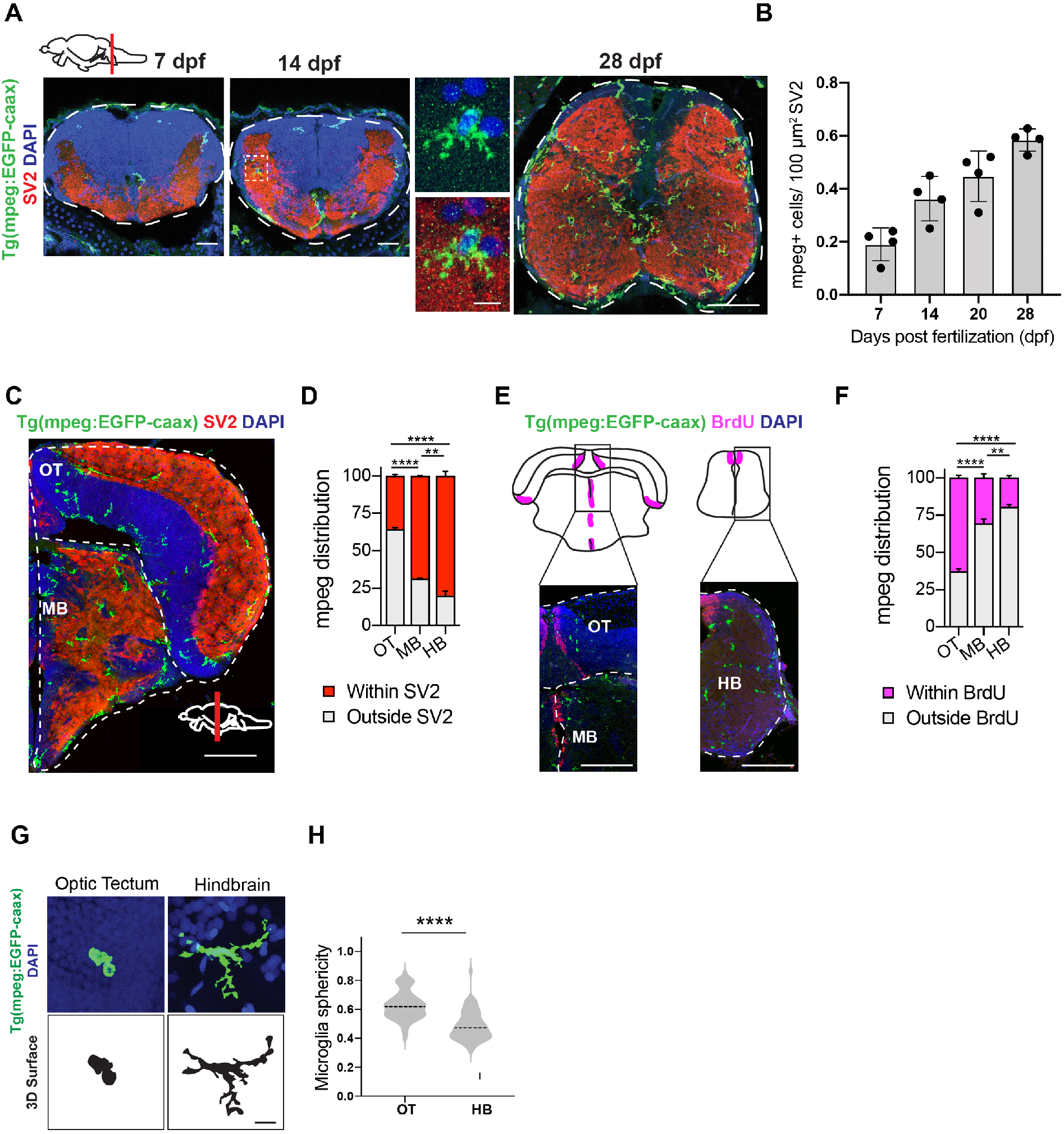
Phenotypically distinct populations of microglia in synaptic and neurogenic regions of zebrafish brain. **(A)** Representative images of microglia (mpeg:EGFP) and synapses (SV2 antibody stain) in developing zebrafish hindbrain. Inset: representative synapse-embedded microglia at 14 dpf. Scales: 7-14 dpf, 20μm; 28 dpf, 50μm; inset, 5μm. **(B)** Quantification of mpeg:EGFP+ cells per 100 μm^2^ of SV2+ synaptic area at 7,14, and 28 days post fertilization. Dots represent individual fish, data are mean ± SD. **(C-D)** Representative images and quantification of the proportion of mpeg:EGFP+ cells in synaptic (SV2+; red) vs. cellular (DAPI+/SV2-; grey) areas. Distribution quantified within each brain region as outlined (dotted lines): midbrain (MB), optic tectum (OT), and hindbrain (HB, see panel 1A). Mean ±SEM from 4 fish. Scale: 50μm. **(E-F)** Representative images and quantification of the proportion of mpeg:EGFP+ cells within 20 μm of BrdU+ neurogenic regions (purple) vs. outside neurogenic regions (grey). Mean ±SEM from 3 fish. Scale: 50μm. **(G)** Representative images of mpeg:EGFP+ microglia and thresholded maximal projections in OT and HB at 28 dpf. Scale: 5μm. **(H)** Quantification of microglial sphericity (Scale 0-1, 1=most spherical/ameboid), from images thresholded as in G. n= 50 microglia from n=4 fish. *Statistics:* Two way ANOVA with Tukey’s post-hoc comparison (D,F), two-tailed unpaired t-test (H, K). *p < 0.05, **p<0.01, ***p<0.001, ****p,0.0001 **See also Fig S1.**

### Molecularity distinct subsets of microglia identified at single cell resolution during brain development

To determine whether microglia in the zebrafish brain are molecularly heterogeneous, we performed single-cell RNA sequencing. We flow sorted hematopoietic cells from juveniles at 28 dpf using a CD45:DsRed reporter ^37^ crossed with myeloid-specific mpeg:eGFP to ensure that all potential subsets of immune cells were captured (**Fig. 2A, S2A-B**). Our juvenile time-point at 28 dpf encompasses both developmental waves of embryonic and adult microglia in the zebrafish brain ^29,30^. Flow analysis and our initial single-cell clustering indicated that 90% of CD45+ cells were mpeg-EGFP positive, and that the CD45+ population captured all mpeg:eGFP positive cells. Unbiased clustering of CD45+ cells revealed 15 distinct clusters of hematopoietic origin, including seven non-myeloid clusters (*mpeg^neg^*;**Fig. 2B, S2B-C; Table S1**). These included clusters that expressed markers for T-cells (*cd4-1, lck*, and *ccr9a*), natural killer cells (*eomesa*), and innate lymphocyte-like cells (*il13, gata3*) ^38–41^; **Fig. S2C**). Thus, multiple immune cell subsets, predominately myeloid in origin, are present in the zebrafish brain, although it is possible that some of these may be circulating rather than tissue-resident.

**Figure 2.**
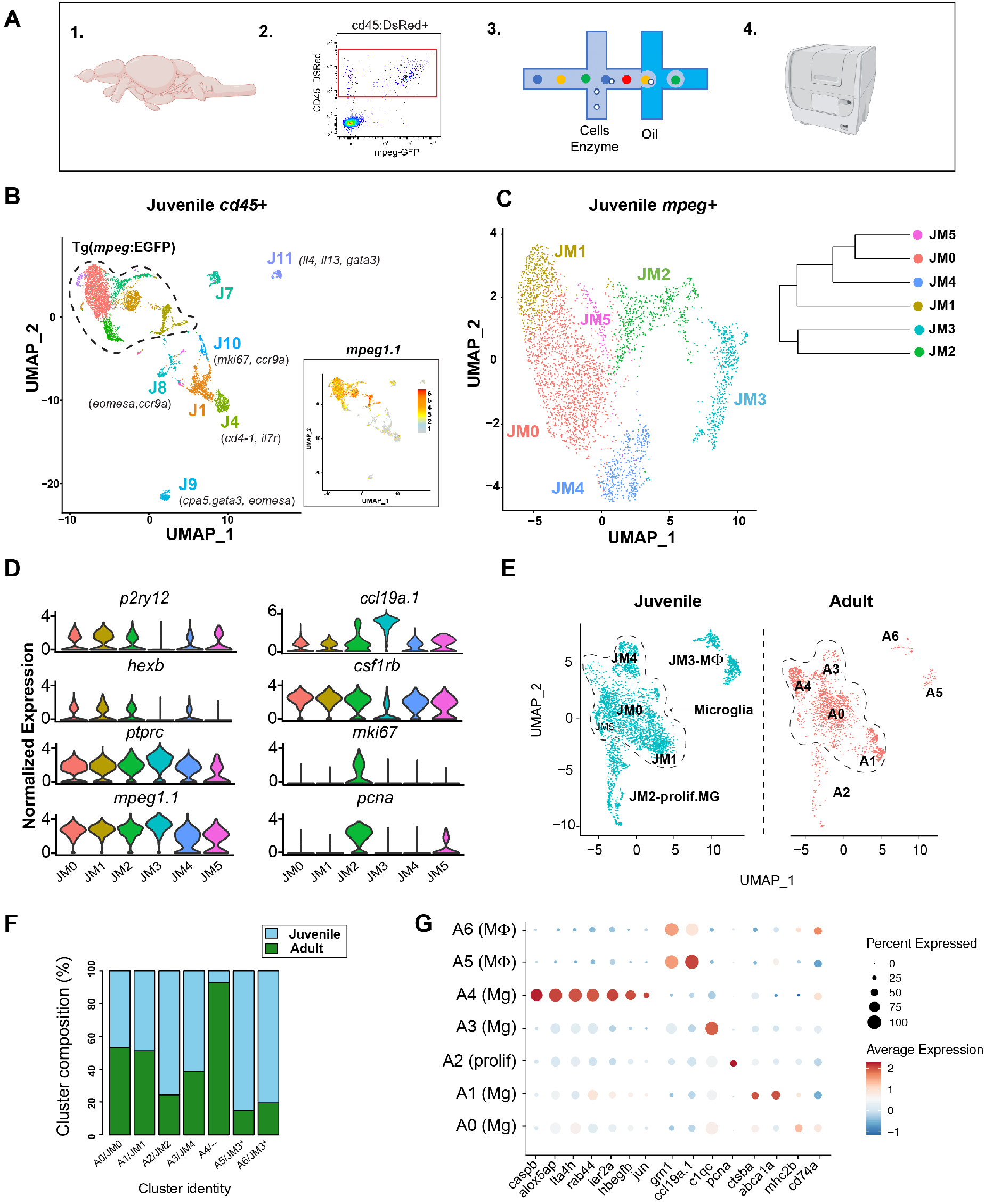
Molecularity distinct subsets of microglia identified at single cell resolution during brain development and in adulthood. **(A)** Schematic diagram of whole brain scRNA -sequencing pipeline. **(B)** Unsupervised clustering of 6,666 juvenile *ptprc (CD45)+* cells. Inset: feature plot of *mpeg1.1* expression. n= 10 fish in two independent replicates. **(C)** Subclustering of 3,539 *mpeg1.1+* cells in C. Inset: cluster dendrogram. **(D)** Violin plots for select marker genes across juvenile clusters from C (JM1-5), including microglia (*p2ry12, hexb*, and *csf1rb*), pan-myeloid (*mpeg1.1*), pan-hematopoietic (*ptprc/cd45*), macrophage (*ccl19a.1*) and proliferative (*mki67* and *pcna*). **(E)** UMAP plots showing co-clustering analysis of juvenile cells in C and 2,080 adult myeloid (*mpeg1.1 +*) cells. Conserved macrophage and proliferative clusters indicated. Dotted line outlines remaining ‘homeostatic’ microglial clusters. **(F)** Percent of each cluster shown in E consisting of Juvenile (blue) vs. Adult (green) derived cells. Normalized within groups to account for overall cell number difference between ages. **(G)** Dot plot highlighting select up- and downregulated genes within adult clustering in E. **See also Fig S2.**

To focus on myeloid cells, which are the dominant immune cell subset, we reclustered 3539 mpeg+ cells, which after quality control and filtering, yielded six distinct myeloid clusters (**Fig. 2C; Fig. S2D-F; Table S2**). We identified cluster JM3 as the macrophage subset based on the absence of microglial-specific markers (*p2ry12, csf1rb, hexb, and slc7a7*) and the presence of *ccl19a. 1 (a.k.a*. macrophage inflammatory protein; **Fig. 2D**) and higher levels of mpeg:EGFP; **Fig. S2G**). Cluster JM2 marked a proliferating subset (*pcna* and *mki67;* **Fig. 2D**). Cluster JM5 contained too few cells to be definitively assigned. The remaining three clusters (JM1, JM0, JM4) had microglial identity but distinct gene expression profiles.

Next, we examined whether these subsets persist into adulthood and mapped their regional distribution in the brain by co-clustering our juvenile data with adult brain microglia that were sorted and sequenced in parallel. The Harmony R package ^42^ was used to integrate the two data sets and compensate for sources of technical variation. Unbiased clustering and differential expression analysis revealed six groups that were conserved between juveniles and adults (**Fig. 2E; Table S3**) as well as a distinct adult-enriched microglial cluster (A4; **Fig. 2F**). Notably, adult cluster A3 mapped to juvenile cluster JM4 (conserved genes included *cebpb, c1qa, c1qb*, and *c1qc*), whereas adult cluster A1 mapped to juvenile cluster JM1 (*apoeb* and *ctsba*). A proliferative cluster was still present (A2; *pcna, tubb2b*, and *mki67*), as well as two distinct macrophage clusters (A5 and A6) corresponding to juvenile cluster JM3 (*ccl19a, siglec15l*, and *cmklr1*). Interestingly, the novel adult-enriched cluster A4 was distinctly enriched in inflammasome genes, suggesting a microglial subset poised for inflammasome activation (*caspb, fads2, and alox5ap;* **Fig. 2G**). Taken together, these data reveal myeloid cell heterogeneity in the juvenile zebrafish brain that persists in the adult.

### Region-specific transcriptional signatures of juvenile zebrafish myeloid cells

Previous zebrafish transcriptomes using the Tg(*mpeg*:EGFP) transgenic line and bulk sequencing approaches have suggested regional transcriptional heterogeneity within the zebrafish brain ^31^. To better compare our single cell data with the existing literature, we isolated *mpeg:EGFP+* myeloid cells from OT, midbrain, and hindbrain at 28 dpf by flow cytometry and performed bulk RNA sequencing (**Fig. 3A, Fig. S3A-B, Table S4**). The resulting transcriptome was highly enriched for microglial specific genes (*csf1rb, c1qa*, and *p2ry12*), as well as some macrophage markers (*siglec15l, spock3*). This was a pure myeloid population with no evidence of neuronal, astrocyte, or oligodendrocyte genes (**Fig. S3C**). Principal component analysis revealed that 60% of gene variance was driven by brain region (PC1; **Fig. 3B**), and differential expression analysis of these subsets (p<0.05) identified region specific gene expression signatures (**Fig. 3C**). In particular, complement and antigen presentation genes were enriched in the hindbrain (*c1qa, c1qc, cd74a*, and *mhc2dab*), whereas lysosome-associated genes were highly enriched in the OT (*apoeb, ctsba, ctsc*, and *ctsla*). The midbrain was intermediate and did not clearly segregate with either phenotype.

**Figure 3.**
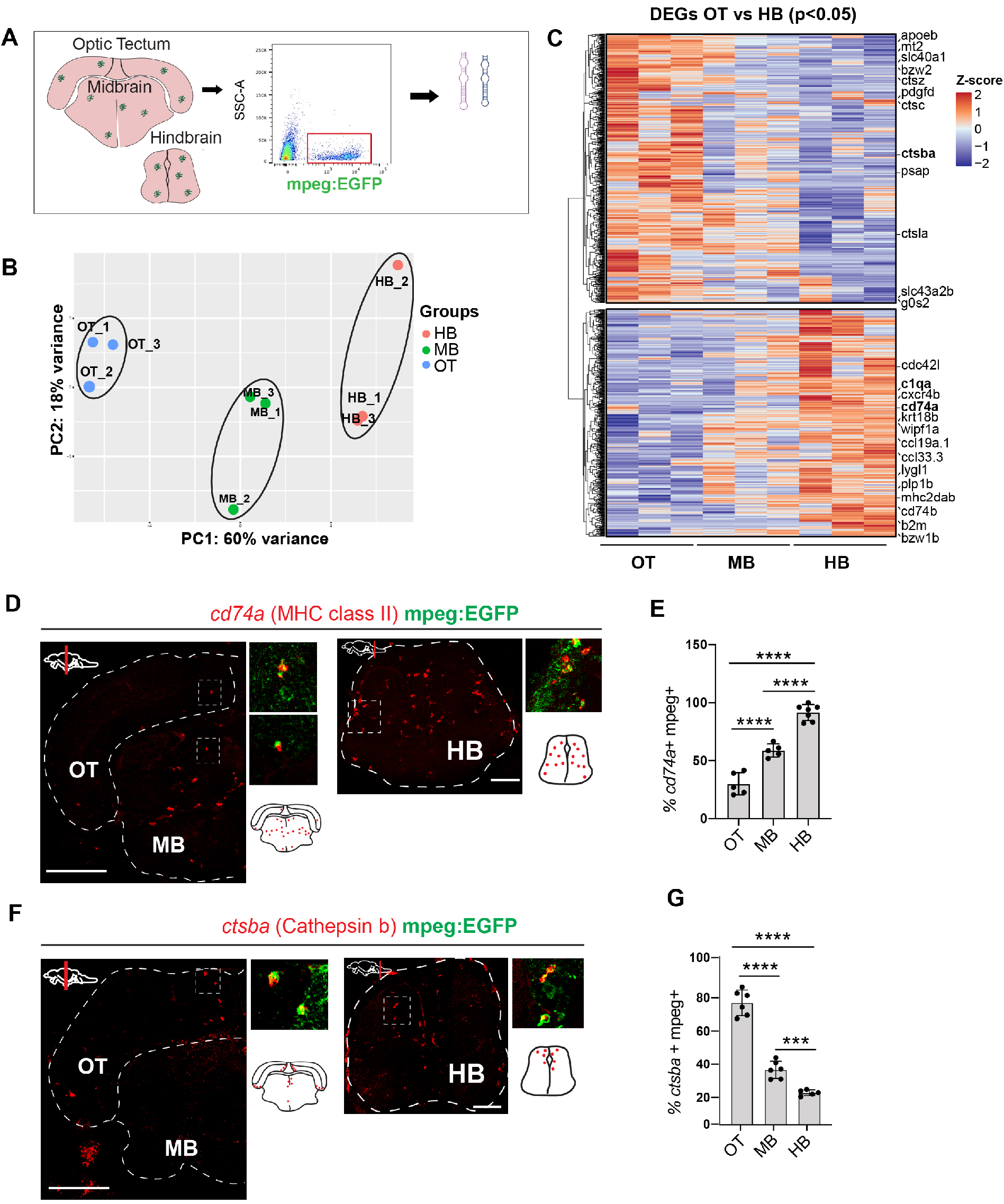
Region specific transcriptional signatures of juvenile zebrafish myeloid cells. **(A)** Schematic of region-specific bulk-sequencing approach using *mpeg*:EGFP. **(B)** Principal component analysis (PCA) plot of top 500 most variable genes for OT, MB, HB regions (dots= independent replicates from 10 pooled fish/replicate). **(C)** Heatmap of differentially expressed genes (DEGs) from mpeg:EGFP positive cells across brain regions (Adjusted p-value <0.05). Selected top differentially expressed genes from each cluster highlighted. **(D-E)** Representative images and quantification of *cd74a* expression across brain regions by *in situ* hybridization (ISH), co-localized with mpeg-GFP. Quantification shows percentage of cd74 + mpeg+ microglia within each brain region. Scale: 50μm. **(F-G)** Representative images and quantification of *ctsba* expression by ISH colocalized with mpeg-GFP. Quantification shows percentage *ctsba*+ mpeg+ microglia within each brain region. Scale: 50 μm. *Statistics:* In E,G, One way ANOVA with Tukey’s post hoc test. Mean±SD, dots=individual fish. *p < 0.05, **p<0.01, ***p<0.001, ****p<0.0001. **See also Fig. S3.**

We confirmed region-specific expression for two candidate genes from this profile via *in situ* hybridization (ISH). The HB enriched gene *cd74a* was expressed in over 90% of hindbrain microglia (mpeg+4c4+) and 30% of OT microglia, and intermediate in the midbrain (**Fig. 3D-E**). In contrast, the OT-enriched gene *ctsba* was highly enriched in neurogenic areas and highest in the OT compared to midbrain and hindbrain (72%, 33%, 19%, respectively; **Fig. 3F-G**). These data reveal brain-region specific transcriptional signatures that can be identified *in situ*. However, they also suggest that these differences may be partly driven by varying proportions of phenotypically distinct microglia as well as resident macrophages that are better discriminated at the single-cell level of resolution.

### Region-enriched functional microglial subsets in zebrafish hindbrain and optic tectum

We next used our region-specific transcriptional signatures to map our single cell data, with the goal of localizing functional subsets *in situ* (**Fig. 4A**). To do this, we calculated an “eigengene” composed of the top differentially expressed genes in the mpeg+ population from each region. Overlay of these region-defining eigengenes onto our UMAP plot revealed substantial enrichment of the OT signature in the juvenile cluster JM1, whereas the hindbrain eigengene was enriched in JM4 and the macrophage cluster JM3 (**Fig. 4B-C).** The midbrain regional signature was indeterminate and aligned only with the macrophage cluster, consistent with our finding of substantially more macrophages in this region **(Fig. S1D**). Cluster JM0 was not enriched in any regional signature, suggesting that it may represent a common microglial population, or a subset not represented in our bulk sequencing. Since the OT-enriched cluster JM1 and the hindbrain-enriched cluster JM4 were distinct in both bulk and single-cell sequencing and did not contain contaminating macrophages, we directly compared the transcriptomic profile of these subsets **(Fig. 4D-E).**

**Figure 4.**
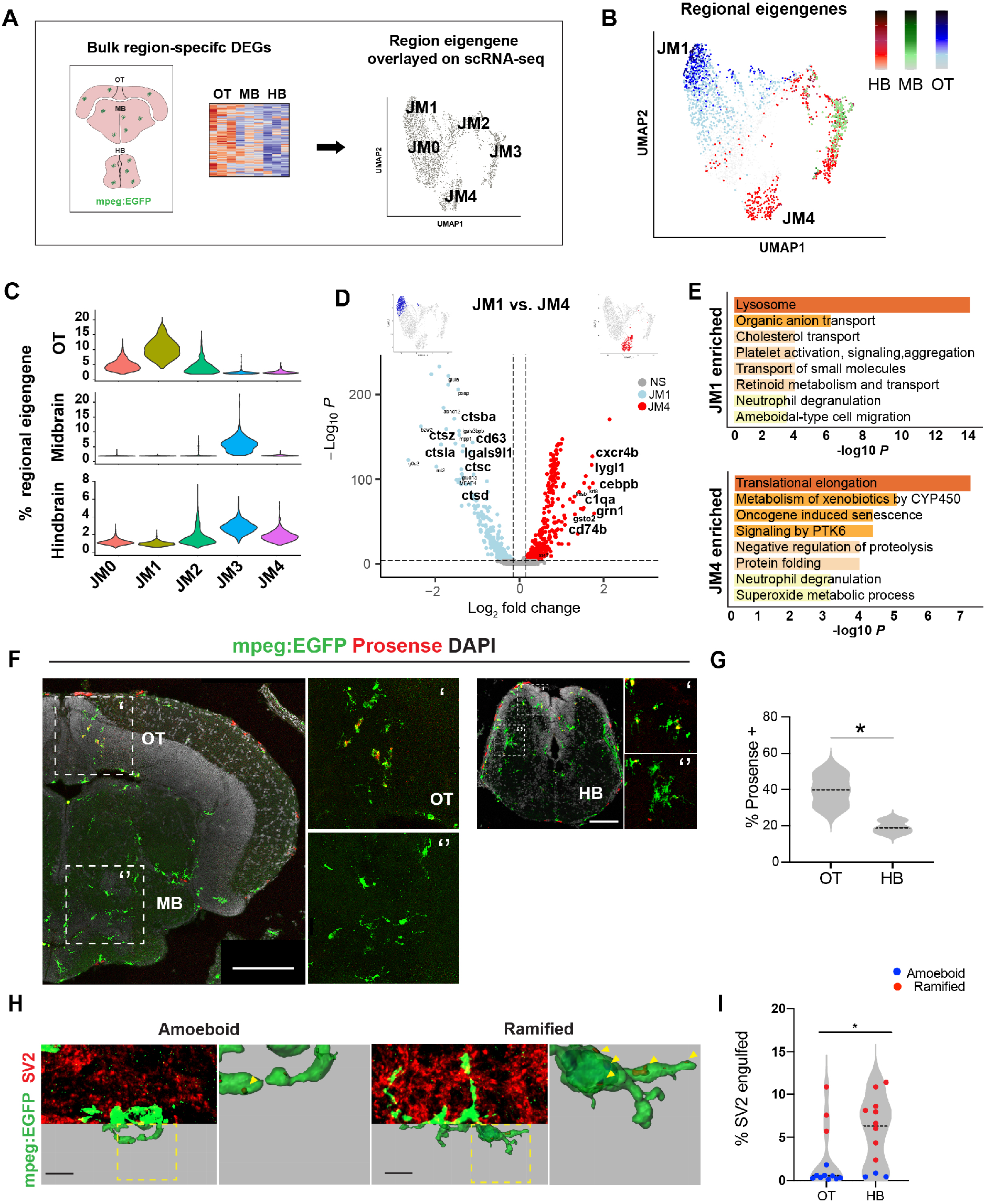
Region-enriched functional microglial subsets in zebrafish hindbrain and optic tectum. **(A)** Schematic of analysis pipeline overlaying brain-region defining genes identified with bulk sequencing onto mpeg+ scRNA seq clusters. **(B)** Feature plots of region-defining eigengenes from hindbrain (HB; red), midbrain (MB; green) and optic tectum (OT; blue) overlaid on mpeg+ UMAP, highlighting microglial clusters JM1 and JM4. **(C)** Violin plots of regional eigengene distributions, related to B. **(D)** Volcano plot of differentially expressed genes between clusters JM1 (OT-enriched; blue) and JM4 (hindbrain-enriched; red). Thresholds represented by dotted lines were set to adjusted p value <10^-8^, log fold change > 0.2. **(E)** Top GO terms from differentially expressed genes in D. **(F)** Representative images of cathepsin-cleaved Prosense 680 colocalized with mpeg:EGFP in indicated brain regions. Scale: 100 μm. **(G)** Quantification of percent microglia containing cleaved Prosense680 in OT and HB. Unpaired t-tests. Mean ± SD. *p < 0.05. OT (Optic Tectum) and HB (Hindbrain). **(H)** Representative images and 3D reconstructions of mpeg:EGFP+ microglia with engulfed SV2 protein. Insets: close-up of reconstructions (arrowheads:SV2 content). Scale bar 5 μm. **(I)** Quantification of percent microglial volume containing SV2 in randomly selected microglia from OT and HB. Post hoc analyses show morphology assignment as ramified (red, sphericity<0.6) vs. amoeboid (blue, sphericity >0.6). Unpaired t-tests within regions, ***p < 0.001, dots= mean value per fish from 3 microglia per fish. *Statistics:* Two-tailed unpaired t-test (G, I). *p < 0.05, **p<0.01, ***p<0.001, ****p,0.0001

Genes that defined the hindbrain-enriched cluster (JM4) included *c1qa*, the initiating protein in the complement cascade, a known regulator of synaptic engulfment expressed in rodent microglia ^43,44^. Another was *cebpb*, a transcription factor that promotes microglial homeostasis in neurodegenerative disease^45^. Other genes included the hindbrain region defining gene *cd74b*, which encodes the invariant chain of major histocompatibility complex class II, and *grn1*, which encodes progranulin and interacts with C1q to regulate synaptic pruning ^46^. We also identified several putative functional genes not previously studied such as the lysozyme gene *lygl1*. Gene ontology analysis suggested that the preferentially expressed genes are involved in protein production and metabolism, including translational elongation, negative regulation of proteolysis, and protein folding (**Fig. 4D-E**). In contrast OT-enriched cluster JM1 was highly enriched for lysosomal activity, including multiple genes for lysosomal proteases including cathepsins (*ctsba, ctsz, ctsla, ctsc*, and *ctsd*) as well as higher levels of *apoeb* and *lgals9l1*. Thus, functional subsets identified by scRNA seq correlate with the differentially enriched populations of synapse-associated and neurogenic associated microglia that we identified *in situ*.

We hypothesized that hindbrain-enriched cluster JM4 represents a synapse-associated microglia subset whereas JM1 preferentially associates with neurogenic regions. To identify our putative OT cathepsin-rich cluster (JM1) *in situ*, we quantified functional cathepsin activity with the biomarker Prosense 680, which becomes fluorescent after proteolytic cleavage by cathepsins with a preference for cathepsin B and L ^47^. Microglial cathepsin activity (mpeg+prosense+) was highly enriched in the OT, where it was mostly detected in ameboid microglia around neurogenic regions (**Fig. 4F-G**), closely matching the expression of *ctsba* by *in situ* hybridization (**Fig. 3F-G**). Consistent with this, tectal microglia engulfed substantially more cell corpses, as quantified by uptake of a neuronal-soma enriched DsRed *Tg(NBT:DsRed)* compared to the hindbrain (**Figure S4A-B**). Next, we quantified synapse engulfment by 3D reconstruction of mpeg-EGFP positive cells with the synaptic marker SV2. We found that hindbrain microglia engulfed significantly more SV2 than OT microglia, consistent with enrichment of the JM4 cluster in hindbrain (**Fig. 4H-I**). Importantly, stratifying this data by morphology revealed that ramified microglia (sphericity ¼0.6) engulfed more SV2 regardless of brain region, although they were markedly more abundant in the hindbrain. These data suggest functional subsets whose abundance differs between brain regions, rather than strictly region-specific functions. Taken together, we conclude that our cathepsin-enriched scRNA Seq cluster JM1 corresponds to cell corpse-engulfing microglia highly enriched in neurogenic regions, whereas complement-expressing cluster JM4 identified a synapse-associated subset.

## Discussion

Here we define the regional localization and molecular signatures of two distinct microglial phagocytic states in the developing zebrafish, including a novel subset of synapse-associated microglia (SAMs) abundant in the hindbrain (**Fig. 5 A, B**). This functionally annotated single cell dataset is an opportunity to investigate fundamental questions related to interactions between microglia and synapses. For example, microglia both engulf synapses and promote synapse formation via modifying the extracellular matrix and other mechanisms ^1,48,49^ The molecular mechanisms that regulate these different states are not well understood. The impact of neuronal activity on microglial function is also a major area of interest: microglial synapse engulfment has been proposed to be activity-dependent, and in both fish and rodents microglial contact can acutely regulate neuronal activity ^50,51^. However, observing these processes in the intact developing brain is extremely challenging in rodents, and is a major strength of the zebrafish model. This molecularly defined population of synapse-associated microglia in zebrafish hindbrain provide an opportunity to temporally define and genetically manipulate these microglial subsets in physiological and in disease contexts.

**Figure 5.**
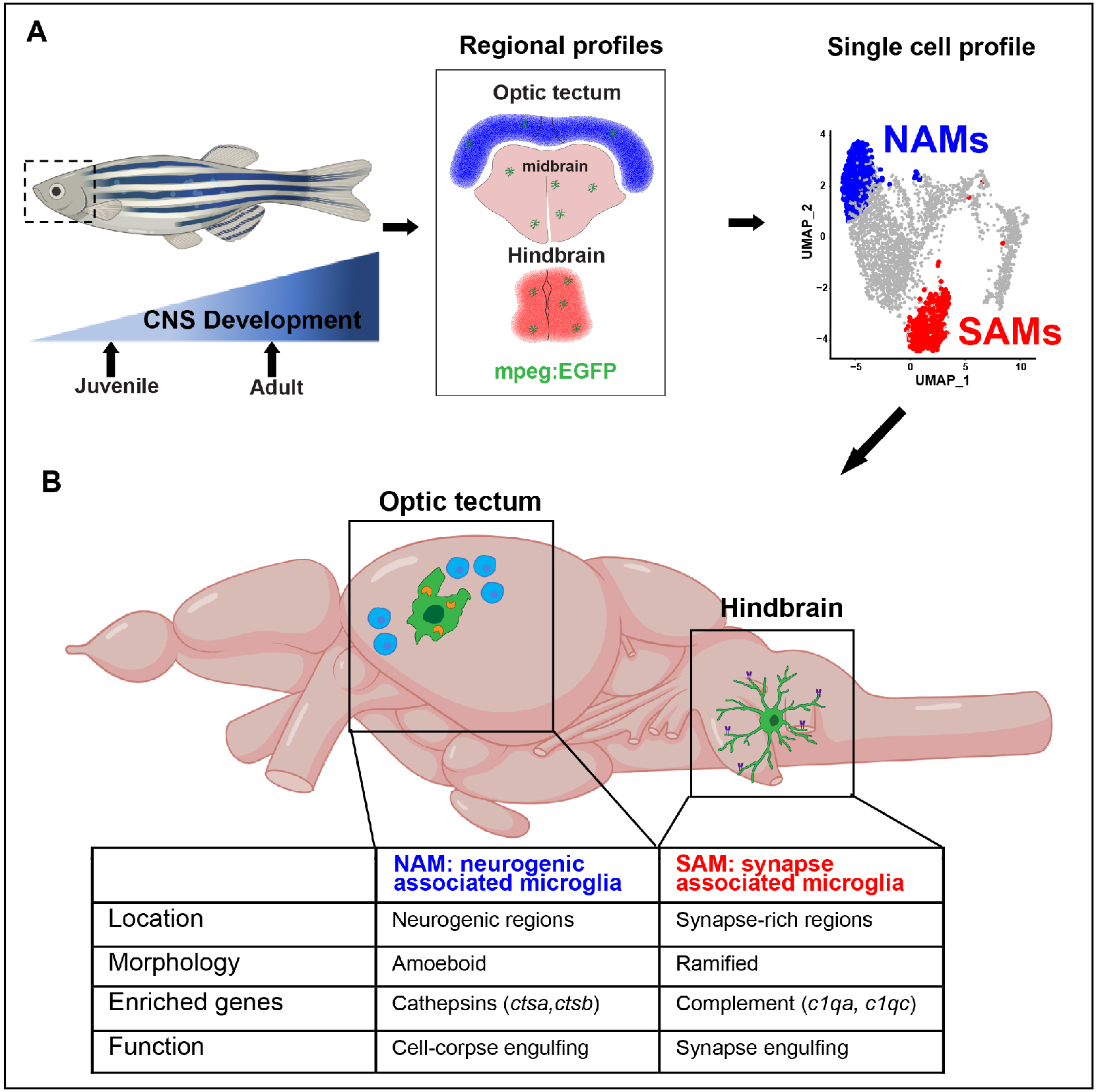
Summary diagram of the identified neurogenic-associated microglia (NAMs) and synapse-associated microglia (SAMs) in the developing zebrafish brain. **(A)** Schematic of analysis pipeline overlaying brain-region defining genes identified with bulk sequencing onto mpeg+ scRNA seq clusters identifies NAMs and SAMs. **(B)** Unique features of NAMs and SAMs in the developing juvenile brain.

Importantly, we show that both SAMs and NAMs are phagocytic *in vivo*, but in different contexts and via distinct molecular mechanisms. Our observations regarding NAMs are consistent with several recent publications: deficits in lysosome function result in defective cell corpse phagocytosis and dysmorphic ‘bubble microglia’ in the OT ^20^. Another study used bulk RNA sequencing to identify *ccl34* as a marker of these ameboid phagocytic microglia in the adult zebrafish, a marker which is also significantly enriched in our NAM subset ^31^. Our study defines the transcriptome of this subset at single cell resolution, and establishes an in vivo assay to track lysosomal function in this population. More importantly, we have identified novel genes associated with a microglial population that is not well-described in the zebrafish -- synapse-associated microglia. The initiating components of the classical complement cascade (*c1qa* and *c1qc*) are top differentially expressed genes in our SAM subset. C1q is also microglial-encoded in the murine brain and in multiple vertebrate species including zebrafish ^33,43^ and promotes developmental synaptic engulfment ^52^. Other top candidates, including the transcription factor *cebpb* and major histocompatibility complex, class II (*cd74a & cd74b*), have been implicated in microglial function in rodent models of Alzheimer’s disease ^45,53^. These data suggest that the SAM profile identifies a core microglial program conserved across species, and raise the question of how other gene candidates, including *lygl1, grn1, fgl2a* and others, are regulating microglial-synaptic interactions.

Our data highlight evolutionarily conserved features of zebrafish microglia as well as unique strengths of the zebrafish model. It is striking that developmental microglial heterogeneity in the fish is largely conserved into adulthood, in contrast to generally diminishing amounts of diversity in adult rodents ^3,4^ This may reflect the fact that fish continue to grow in size throughout life, adding new neurons and new synaptic connections. This suggests that the time window for studying microglial roles in circuit remodeling may be much broader than in mammals, and that mechanisms may exist to maintain synaptic plasticity throughout life. Future studies focusing on microglial-synapse interactions in the zebrafish hindbrain can take advantage of co-existing populations of cell-corpse engulfing and synapse associated microglia to further define the differences between these subsets. Our data also suggest that live-imaging of microglial-synapse interactions may help to answer key questions about how and why microglia interact with synapses. Finally, the ability to do high throughput screening in zebrafish raises the possibility that this model could be used to define new therapeutic targets, in neurodevelopmental diseases linked to immune dysfunction including autism spectrum disorder, epilepsy, and schizophrenia.

## Acknowledgements

We are grateful to Marci Rosenberg, Haruna Nakajo, and members of the Molofsky Lab for helpful comments on the manuscript, to Ari Molofsky, Tom Nowakowski and Galina Schmunk for their expert advice, and to Brian Black, Gary Moulder and Louie Ramos for the support of the CVRI Zebrafish Core facility. Thanks to Francesca Peri for the *Tg*(*mpeg:EGFP-caax*), David Traver for the *Tg(Cd45:DsRed)* fish, and Roland Wu for the *Tg(mpeg:EGFP)* fish, and to the UCSF Laboratory for Cell Analysis and Dr. Eric Chow at Center for Advanced Technologies for technical contributions.

## Author contributions

Conceptualization: N.J.S, A.V.M, I.D.V., and L.C.D.; Methodology, N.J.S., I.D.V, and L.C.D; Investigation, N.J.S, I.D.V, L.C.D, and N.C.H; Writing – Original Draft, N.J.S, L.C.D., and A.V.M.; Writing – Review & Editing, all co-authors; Funding Acquisition, A.V.M. and N.J.S. Resources, A.V.M.; Supervision, A.V.M. and I.D.V

## Data and materials availability

Supplement contains additional data. All data needed to evaluate the conclusions in the paper are present in the paper or the Supplementary Materials. Single cell RNA sequencing and bulk RNA sequencing data were deposited in NCBI’s Gene Expression Omnibus (Barrett et al., 2013) under accession numbers GSE164772 and GSE164771, respectively.

## Supplemental information

**Figs S1-S4**

**Tables S1-S3**

**Methods**

---

**Figure S1.**
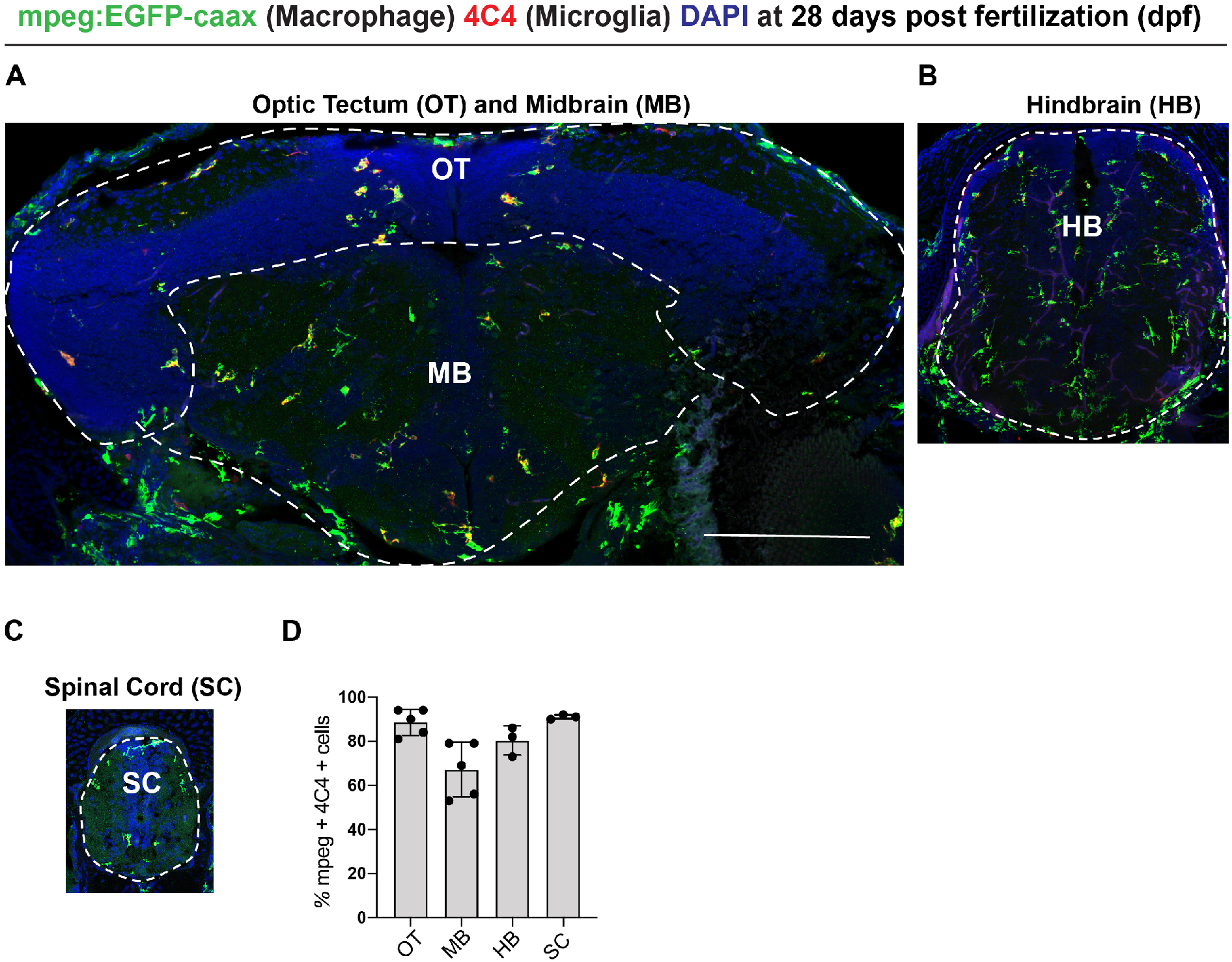
Macrophages and microglia in OT, MB, and HB regions at 28 dpf, supplemental to Figure 1. **(A-C)** Representative images of brain regions using the pan-myeloid reporter line mpeg:EGFP and the microglial-specific antibody 4C4 at 28 days post fertilization (dpf). **(D)** Quantification of percent of mpeg:EGFP+ population also expressing 4C4. Scale: 100 μm. Optic tectum (OT), midbrain (MB), and hindbrain (HB).

**Figure S2.**
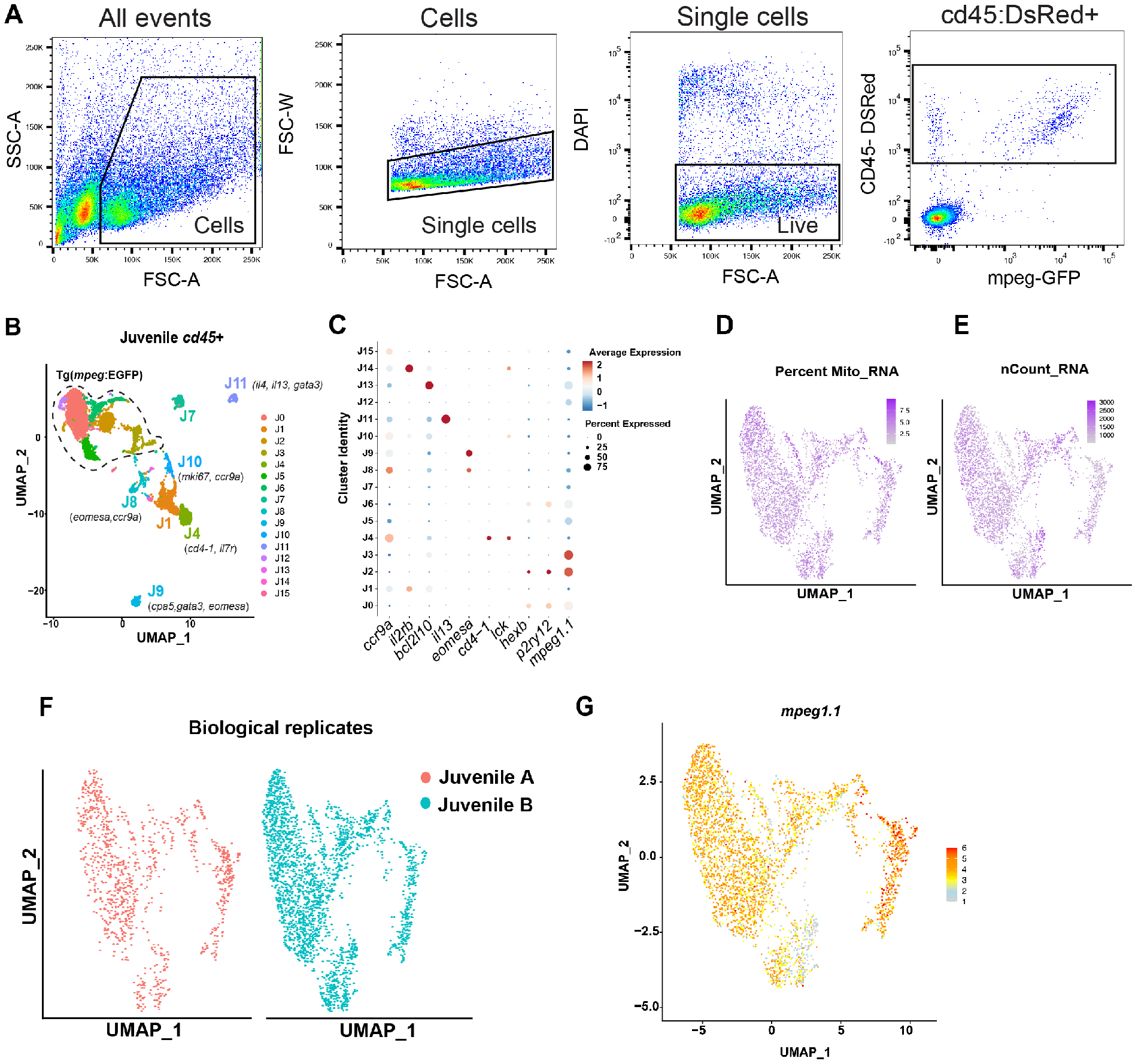
Gating hierarchy and quality control of single cell sequencing data, supplemental to Figure 2. **(A)** Gating strategy to isolate *cd45* positive cells for single-cell RNA sequencing. **(B)** Unsupervised clustering of juvenile *cd45* positive cells, replicated from Fig. 2B for reference. **(C)** Select enriched genes in *cd45+mpeg*- immune cell subsets. **(D-E)** Feature plot showing distribution of mitochondrial RNA content and genes recovered (nCount) per cell. **(F)** Comparison of independent biological replicates of mpeg+ cells pooled in Fig. 2C. **(G)** UMAP plot highlighting levels of *mpeg1.1* expression the mpeg1.1+ population from 2C.

**Figure S3.**
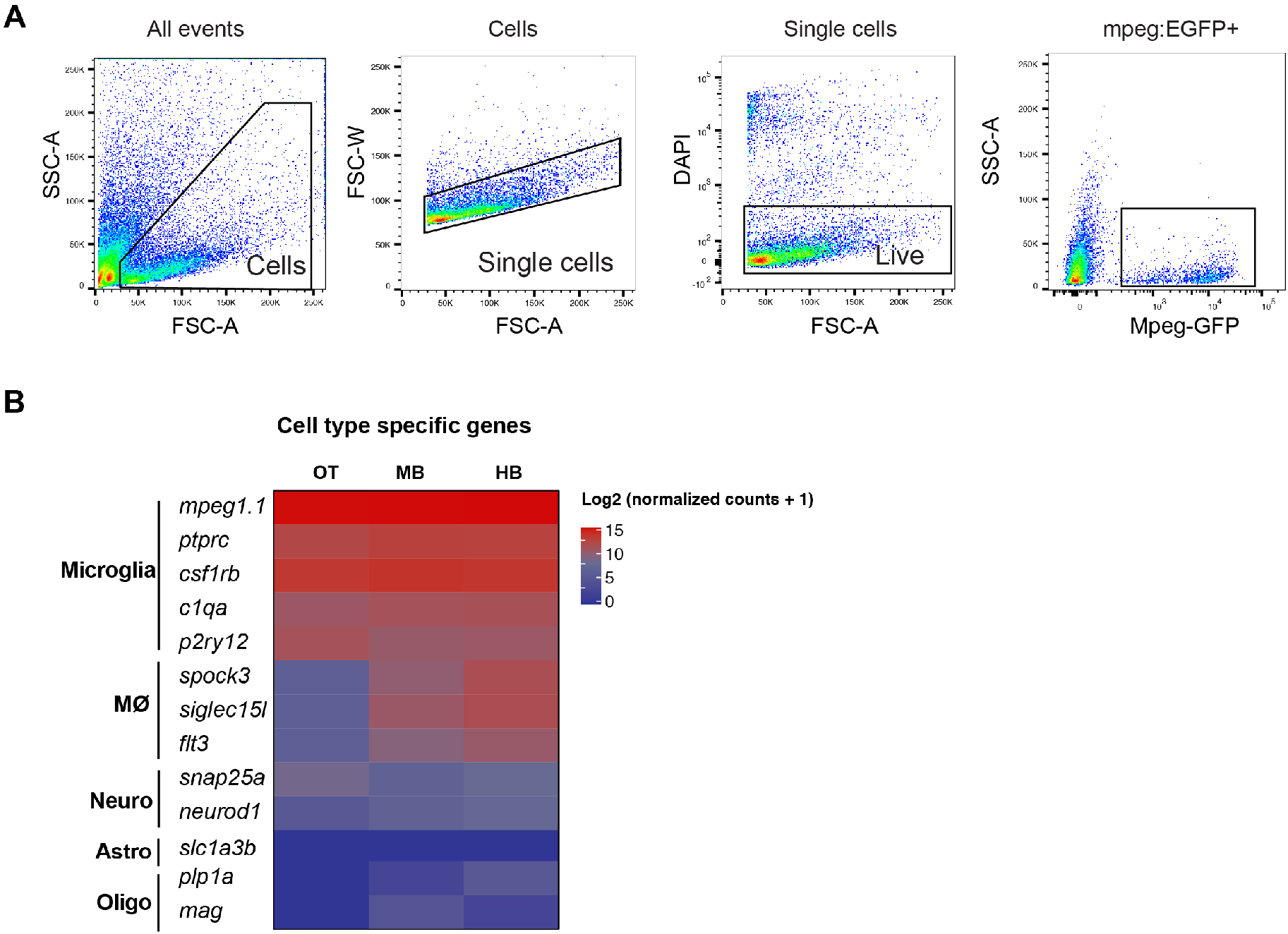
Gating strategy and quality control of bulk sequencing data, supplemental to Figure 3. **(A)** Gating strategy to isolate mpeg-EGFP+ myeloid cells for region-specific bulk sequencing. **(B)** Heatmap of selected microglia, macrophage, neuronal, astrocyte, and oligodendrocyte genes. Optic tectum (OT), midbrain (MB), and hindbrain (HB).

**Figure S4.**
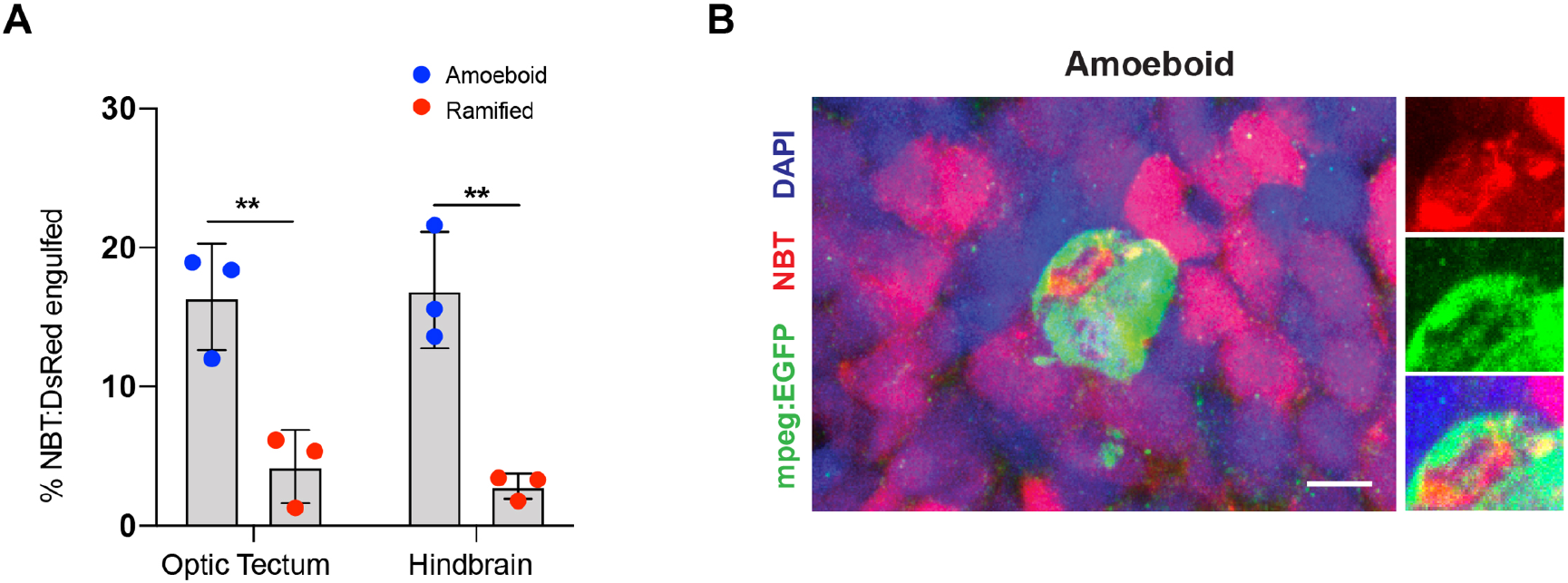
Neuronal engulfment in the OT and HB at 28 dpf. **(A-B)** Representative image and quantification of NBT:DsRed neuronal bodies within ramified and amoeboid microglia (Sphericity > .6) from OT and HB brain regions. Two separate t-test, ***p < 0.001, n=3 microglia per fish represented as mean values (n=3 juvenile fish). Scale bar 10 μm.

## Supplemental excel files

**Table S1: Pan hematopoietic single cell clusters in 28 dpf zebrafish brain**

**Table S2: Myeloid single cell clusters in 28 dpf zebrafish brain**

**Table S3: Adult and juvenile single cell clusters**

**Table S4: Region specific myeloid cell profiling in 28 dpf zebrafish brain**

## KEY RESOURCES TABLE

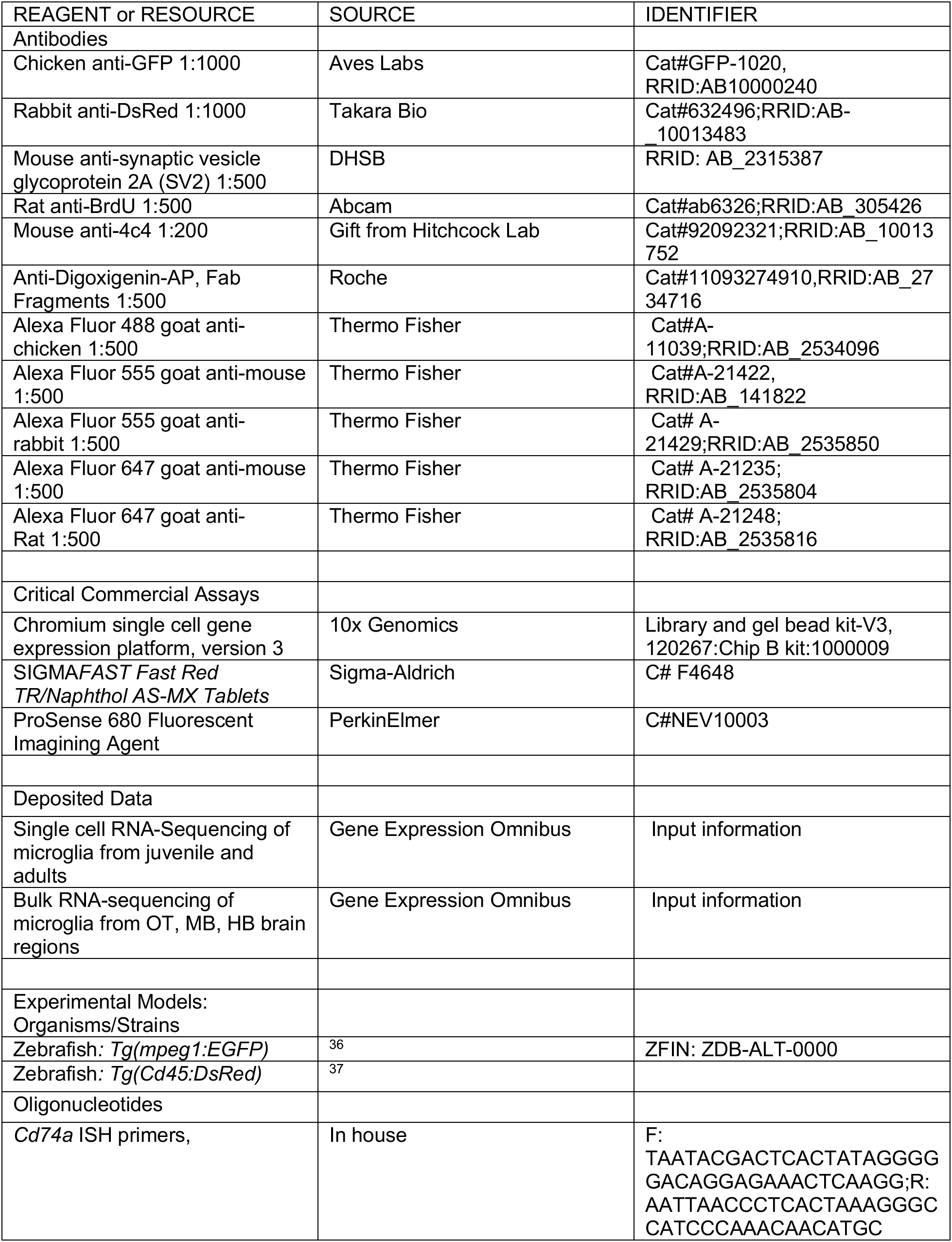
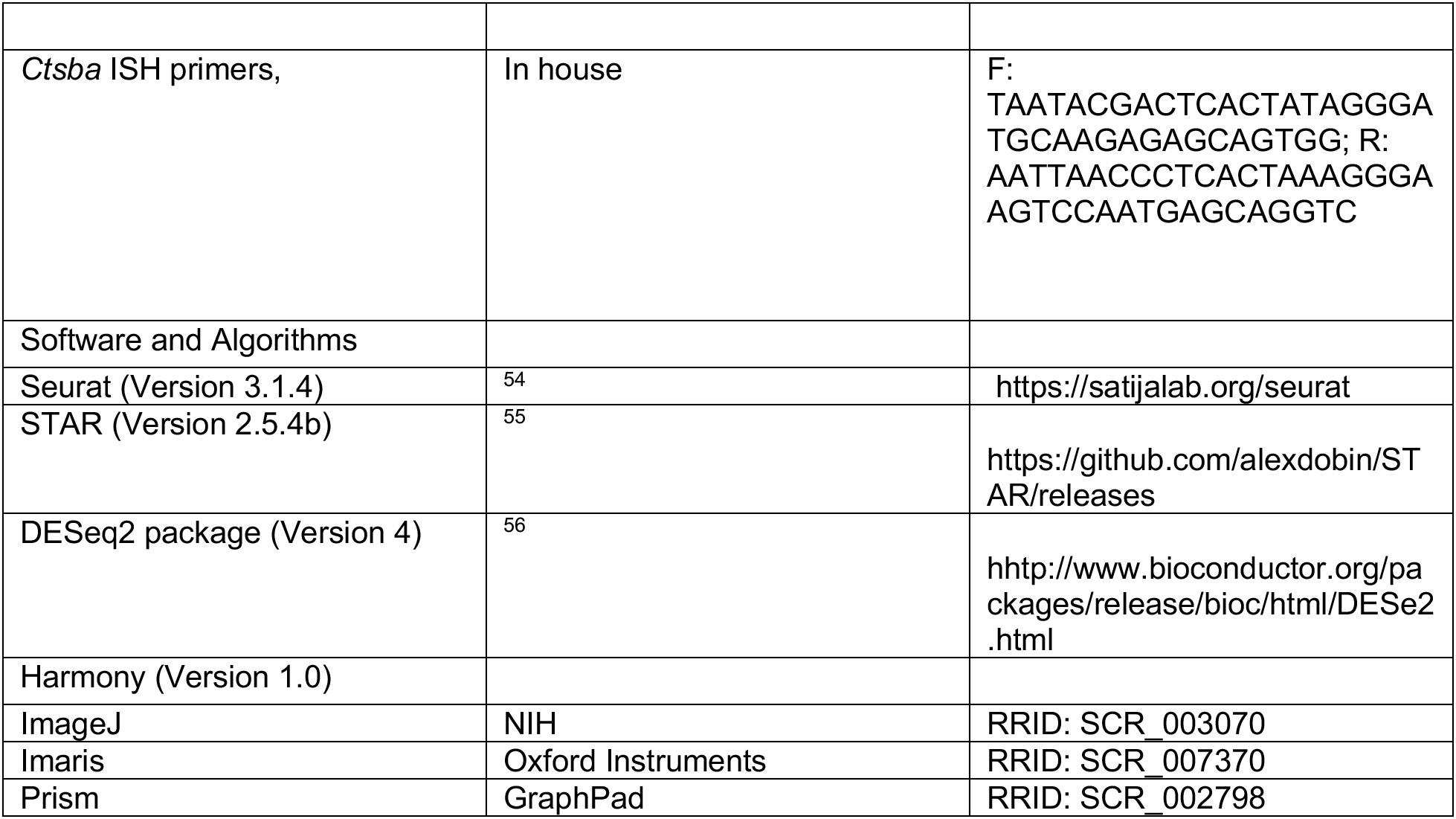

## RESOURCE AVAILABILITY

**Lead Contact** Anna Molofsky

## DATA AND CODE AVAILABILITY

Single cell RNA sequencing and bulk RNA sequencing data were deposited in NCBI’s Gene Expression Omnibus (Barrett et al., 2013) under accession number [GSE164772] single cell and [GSE164771] bulk.

## EXPERIMENTAL MODEL AND SUBJECT DETAILS

Wild-type, AB strain zebrafish (*Danio rerio;* ZIRC, University of Oregon, Eugene, OR) were propagated, maintained, and housed in recirculating habitats at 28.5°C and on a 14/10-h light/dark cycle. Embryos were collected after natural spawns, incubated at 28.5°C and staged by hours post fertilization (hpf). Juveniles used were 28 days of age, a time in development before sex determination. Adults were of either sex were used at 12 months of age. Ages were matched within experiments. The transgenic reporter lines, *Tg[CD45:DsRed]* and *Tg[mpeg1:EGFP]* was used to identify hematopoietic lineage and mononuclear phagocytes ^36,37^. All animal protocols were approved by and in accordance with the guidelines established by the Institutional Animal Care and Use Committee and Laboratory Animal Resource Center.

## METHOD DETAILS

### Immunohistochemistry

Tissues were fixed overnight at 4°C in 0.1M phosphate buffered 4% paraformaldehyde, cryoprotected with 20% sucrose, and embedded in optical cutting temperature (OCT) medium (Sakura Finetek USA, Torrance, CA). Immunohistochemistry (IHC) was performed as previously described (Silva et al., 2020). Briefly, 20-μm-thick sections were collected and mounted onto glass slides. Sections were washed in phosphate buffer saline with 0.5% Triton-x (PBST) and incubated with 20% heat-inactivated normal goat serum in PBST for 2 hours (NSS; Sigma-Aldrich, Corp.). Primary antibodies were applied overnight at 4°C. Sections were then washed with PBST and incubated in secondary antibodies for 1 hour at room temperature. Prior to IHC for BrdU, sections were immersed in 100°C sodium citrate buffer (10 mM sodium citrate, 0.05% Tween 20, pH 6.0) for 30 minutes and cooled at room temperature for 20 minutes. IHC was performed as described above.

### Labeling neurogenic zone

In juveniles, dividing cells were labeled with BrdU by housing animals in system water containing 5 mM BrdU for 24 hours prior to collection ^57^.

### *In situ* Hybridization

*In situ* hybridizations were performed as previously described ^57^. Digoxigenin (DIG)-labeled riboprobes for *cd74a* and *ctsba* was generated by PCR amplification using primers containing the T3 or T7 promoter sequences.

*cd74a* (F): 5’ <u>TAATACGACTCACTATAGGG</u>GGACAGGAGAAACTCAAGG 3’

*cd74a-9* (R): 5’ <u>AATTAACCCTCACTAAAGGG</u>CCATCCCAAACAACATGC 3’.

*ctsba* (F): 5’ <u>TAATACGACTCACTATAGGG</u>ATGCAAGAGAGCAGTGG 3’

*ctsba* (R): 5’ <u>AATTAACCCTCACTAAAGGG</u>AAGTCCAATGAGCAGGTC 3’.

Briefly, 20-μm-thick sections were hybridized with riboprobes at 55°C, incubated with an alkaline-phosphatase-conjugated anti-DIG antibody and visualized using Fast Red TR/Naphthol AS-MX (SIGMA*FAST*) as the enzymatic substrate. When *in situ* hybridizations were combined with BrdU IHC, sections were removed from the fast red solutions, rinsed and post-fixed in buffered 4% paraformaldehyde for 10 minutes then processed for BrdU IHC as described above.

### ProSense680 injections

Juvenile fish were injected with 2 nL of ProSense680 at a concentration of 20 nmol directly into the brain and returned to system water immediately following injections. Fish were euthanized 24 hours post injection and processed for immunohistochemistry as stated above.

### Microglia morphology quantitation

Z-stack images of mpeg:EGFP and 4C4 staining were acquired at the step size of .5μm using a 63x objective (NA 1.4) on an LSM 800 Confocal Microscope (Zeiss) for a thickness of 20 μm. Microglia sphericity was quantified using Imaris software (Bitplane) by creating a 3D surface reconstruction of mpeg:EGFP^+^ microglia that were also positive for 4c4. All images were set to a standard threshold to accurately maintain morphology for quantifications.

### Microglia engulfment assay

Images were acquired with an LSM 800 Confocal Microscope (Zeiss) using the same parameters as described above. Imaris software (Bitplane) was used to generate 3D surface rendering of microglia, which were then masked for NBT:DsRed or SV2 channels within that microglia. Masked channels were then 3D rendered to obtain volume data. NBT:DsRed and SV2 engulfment was calculated per cell as the volume of SV2 divided by the volume of the microglia.

### Fluorescence activated cell sorting (FACS)

For bulk RNA-sequencing of juvenile 28 dpf *Tg(mpeg1:EGFP)* zebrafish, the optic tectum, midbrain and hindbrain were dissected (10 zebrafish were pooled per sample). For scRNA-sequencing of 28 dpf *Tg(mpeg1:EGFP): Tg(Cd45:DsRed)* juveniles (10 zebrafish pooled per lane) and 1 year old *Tg(mpeg1:EGFP): Tg(Cd45:DsRed)* adults (3 zebrafish pooled per lane) whole brains were dissected. To isolate microglia and other CD45+ cells, we followed a previously described method. Briefly, the brain(s) (regions) were mechanically dissociated in isolation medium (1x HBSS, 0.6% glucose, 15 mM HEPES, 1 mM EDTA pH 8.0) using a glass tissue homogenizer (VWR). Subsequently, the cell suspension was filtered through a 70 μm filter (Falcon) and pelleted at 300 g, 4°C for 10 minutes. The pellet was resuspended in 22% Percoll (GE Healthcare) and centrifuged at 900 g, 4°C for 20 minutes (acceleration set to 4 and deceleration set to 1). Afterwards, the myelin free pellet was resuspended in isolation medium that did not contain phenol red. Prior to sorting on a BD FACS Aria III, cell suspension was incubated with DAPI (Sigma). For bulk RNA-sequencing, microglia were gated on FSC/SSC scatter, live cells by DAPI, and Mpeg1-GFP+. After sorting, cells were spun down at 500 g, 4°C for 10 min and the pellet was lysed with RLT+ (Qiagen). For scRNA-sequencing, microglia, macrophages and CD45+ cells were collected by gating on FSC/SSC scatter, live cells by DAPI, and all CD45-DsRed (which included both mpeg+ and negative subsets). After sorting, cells were spun down at 500 g, 4°C for 10 min and resuspended in PBS + 0.05% BSA (Sigma).

### Bulk RNA-sequencing of microglia

RNA was extracted using the RNeasy® Plus Micro kit (Qiagen) from RLT+ lysed microglia. RNA quality and concentration were measured using the Agilent RNA 6000 Pico kit on an Agilent Bioanalyzer. All samples had an RNA Integrity Number (RIN) >8. For each sample, a total of 10 ng of RNA was loaded as input for cDNA amplification and library construction using the QuantSeq 3’ mRNA-Seq Library Prep Kit FWD for Illumina (Lexogen) following manufacturer’s instructions. Library quality was determined with the Agilent High Sensitivity DNA kit on an Agilent Bioanalyzer and concentrations measured with the Qubit^™^ dsDNA HS Assay Kit (Thermo Fisher) on a Qubit^TM^ (Thermo Fisher). Library pools were single-end (65-bp reads) sequenced on two lanes using an Illumina HiSeq 4000 yielding 40-50 million reads per sample.

### Bulk RNA-sequencing analysis

Quality of reads was assessed using FastQC (http://www.bioinformatics.babraham.ac.uk/projects/fastqc). All samples passed quality control and reads were aligned to Danio rerio GRCz11.98 genome (retrieved from Ensemble) using STAR (version 2.5.4b) ^55^ with ‘--outFilterMultimapNmax 1’ to only keep uniquely mapped reads. Uniquely mapped reads were counted using HTSeq ^58^. Subsequently, the DESeq2 package ^56^ in R software was used to normalize the raw counts and perform differential gene expression analysis. Batch correction was done using the Limma package ^59^ and heatmaps were made using ComplexHeatmap ^60^ package in R software. Metascape was used for gene ontology analysis ^61^.

### Single cell RNA-sequencing

Single cells were isolated as described above. Approximately 15,000 cells were loaded into each well of Chromium Chip B (v3), libraries were prepared using the 10x Genomic Chromium 3’ Gene Expression Kit in-house as described in the Manual and sequenced on two lanes of the NovaSeq SP100 sequencer for an average depth of 30,000-50,000 reads per cell. BCL files are converted to Fastq, then used as inputs to the 10X Genomics Cell Ranger pipeline. Samples were processed using the Cell Ranger 2.1 pipeline and aligned to the GRCz11.94 (danRer11) zebrafish reference genome. Clustering and differential expression analysis were conducted using Seurat version 3.1.4. for figure 3 (juvenile) and figure 5 (adult) data.

Feature thresholds

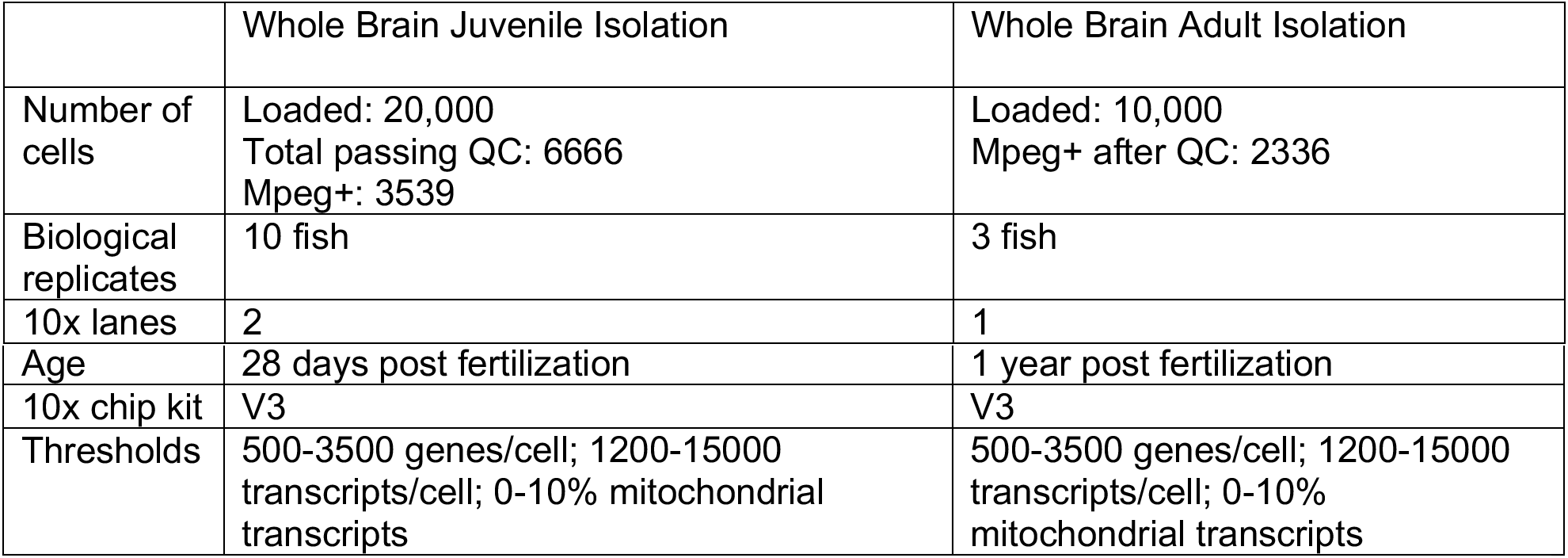

Following alignment in Cell Ranger as described above, counts were imported into R and analyzed using the Seurat package (Butler et al., 2018; Satija et al., 2015). Cells outside of the thresholds listed in the table were excluded from downstream analysis.

Counts were then normalized, regressing out percent mitochondrial RNA and total counts. The top 6000 most variable genes were used to calculate 50 principal components, and the top 30 PCs were used for nearest neighbor, UMAP, and cluster calculations with the resolutions shown in the table. Individual cell types were identified through calculation of marker genes using the MAST test for genes expressed in at least 20% of cells in the cluster and a log fold change of 0.2 or greater and adjusted p value less than 10^-8. The microglial and macrophage clusters were isolated based on expression of CD45 (*ptprc*) and *mpeg1.1*. Normalization, clustering, and differential gene expression was recalculated for each sample (juvenile, juvenile mpeg+, adult and juvenile mpeg+). GO analysis was conducted using the Metascape webpage (www.metascape.org). Adult and juvenile datasets were combined using Harmony 1.0. Bulk vs single cell analysis was conducted using the following thresholds for bulk sequencing results: adjp< 0.05, lfc > 1.2, basemean > 100. Following thresholding, only genes uniquely enriched in one region were used to calculate eigengene values.

## QUANTIFICATION AND STATISTICAL ANALYSIS

Graphpad Prism 8.3.0 was used for all histological quantification analyses. Statistical tests are described in text and figure legends. RNA-sequencing data was analyzed in R as described above.

